# Isotonic tartrazine is sufficient to improve transscleral penetration of optical coherence tomography

**DOI:** 10.1101/2025.11.17.688647

**Authors:** Rahul M. Dhodapkar

## Abstract

Tartrazine dye (FD&C Yellow No. 5) been demonstrated to reduce optical scattering and improve the penetration of optical coherence tomography (OCT) in ex vivo ocular tissues. However, previous reports have used high concentrations of tartrazine that are hyperosmolar to healthy ocular tissues including cornea. Higher concentrations of dye and hyperosmolarity may increase toxicity of tartrazine on the ocular surface, and may limit the potential for clinical translation of these techniques for use in human patients. In this work, we demonstrate that low concentrations of tartrazine dye, approximately isotonic to healthy cornea, are sufficient to produce a measurable improvement in the penetration depth of an OCT scan, while reducing the potential for ocular toxicity.

## 1 Introduction

Since its development in 1991, optical coherence tomography (OCT) has transformed the treatment of vision threatening diseases of the anterior and posterior segments of the eye and has touched nearly every branch of ophthalmology [1]. With modern techniques, OCT can enable the detection of structural changes only 2-3 microns in size [2]. These technologies have been ubiquitously adopted, and OCT findings are central to the management of diverse pathologies such as diabetic macular edema and glaucomatous optic neuropathy [3]. However, OCT’s potential is often limited by the optical properties of the intervening ocular media. It is often impossible to obtain OCT images of the fundus in the setting of corneal scarring or other anterior segment opacifications, and other important structures such as the iridocorneal angle are less clearly visualized behind an optically opaque sclera. The opacity of ocular tissues is due to a combination of light scattering by disorganized collagen fibers in regions such as the sclera, and light absorption by pigment-laden tissues such as the uvea [4].

Recent advancements in optical clearing technologies, especially with potentially biocompatible dyes such as tartrazine (FD&C Yellow No. 5), bring the promise of invivo modulation of light scattering for diagnostic and therapeutic purposes. Tartrazine dye in particular has been shown to enable trans-abdominal optical imaging in live mice at 0.6M concentration [5]. In the eye, tartrazine dye has been shown to improve the transscleral penetration of OCT imaging in porcine tissue using a handheld OCT device at 0.4M concentration [6]. However, the direct application of these techniques for use in humans continues to be limited by concerns regarding potential toxicity.

Tartrazine is an azo dye, and a widely-used food additive that has been extensively studied with regard to its potential toxic effects. While studies are mixed regarding ocular toxicity, at least one report noted that oral administration of tartrazine dye resulted in increased oxidative stress in the eyes of rodents [7]. Azo dyes in general have been described to cause toxicity due to cleavage of azo bonds and the production of reactive oxygen species [8, 9]. While most studies on the toxicity of tartrazine and other azo dyes focus on its oral and systemic absorption, one study found that once-daily topical application of tartrazine 3% w/v (approximately 0.056M) to rabbits for 21 days did not cause significant ocular toxicity or particle embedment [10]. However, the concentration of tartrazine used in previous reports of optical clearing is much higher than what was reported in this ocular topical application study.

In this work, we formulate an isotonic tartrazine aqueous solution and perform OCT penetration measurement of cadaveric unfixed porcine scleral tissue before and after exposure to tartrazine. We show that even at these lower concentrations and shorter exposure times, a measurable improvement in OCT penetration depth was observed. These findings highlight a potential safety benefit from using isotonic tartrazine with short exposure times for in-vivo optical clearing in a clinical context. We believe that these results will provide further support for the use of tartrazine and other clearing dyes as potentially safe agents to improve optical imaging and therapeutics in ocular tissues.

## 2 Methods

Porcine globes were dissected with Westcott scissors and 0.3 forceps to separate the cornea and anterior 4mm skirt of perilimbal sclera from the posterior cup of the globe. Radial incisions were then made in the posterior cup to fashion approximately equal sized scleral samples. Any conjunctiva or Tenon’s, as well as any uveal tissue were sharply removed from the scleral tissue using a Bard-Parker #10 blade, as well as Westcott scissors. Following cleaning of the samples to bare sclera as much as possible, samples were affixed to standard microscope slides using a 6-0 silk suture. All porcine samples were fresh, unfixed, and without any visible signs of tissue degradation at time of use.

To prepare isotonic tartrazine solution, 3 grams of tartrazine powder (534.36 g/mol; Dawn Scientific Category #S10184) were dissolved in 75 mL of sterile water at room temperature to produce a 75 mM solution (300mOsm; 4% w/v). A quenching bath of sterile water was also prepared.

Slide mounted samples were imaged on a Zeiss Cirrus 5000 machine, using the anterior segment OCT settings. Samples were imaged three times sequentially: (1) immediately after preparation, prior to any immersion in tartrazine, (2) after immersion in 75 mM tartrazine for 60 seconds, and (3) after immersion in a quenching bath of sterile water for 60 seconds. OCT penetration depth was manually annotated using measurement tools within the Zeiss Cirrus suite.

Statistical analysis was performed using standard tools from the R programming language [11]. Paired-sample Student’s t-test was used to determine the significant of OCT penetration depth measurements between study time points. Two-tailed tests were used throughout. Data visualization was performed using ggplot2 [12].

## 3 Results

Porcine sclera were dissected from the full globe and cleaned to produce samples of approximately equal size and shape. A single 6-0 silk suture was passed and tied to affix the sample to the glass microscope slide and the samples were found to lay flat in a repeatable way.

Porcine scleral samples immersed in 4% tartrazine solution for 60 seconds showed significantly increased OCT penetration depth as compared to prior to immersion, and after quenching with water (Figure 1). The mean OCT penetration depth prior to tartrazine exposure was 326.4 ± 38.6 µm. After 60 seconds of immersion in 4% tartrazine, the mean penetration depth was 484 ± 44.6 µm. Examination of the gross appearance of the OCT scans was consistent with increased penetration and improved visualization of the superficial layers of the sclera (Figure 1). On unmagnified visual inspection of the samples after immersion, the scleral samples were found to have acquired an orange stain, but the apparent visual transparency of the samples was not noticeably altered.

**Fig. 1.**
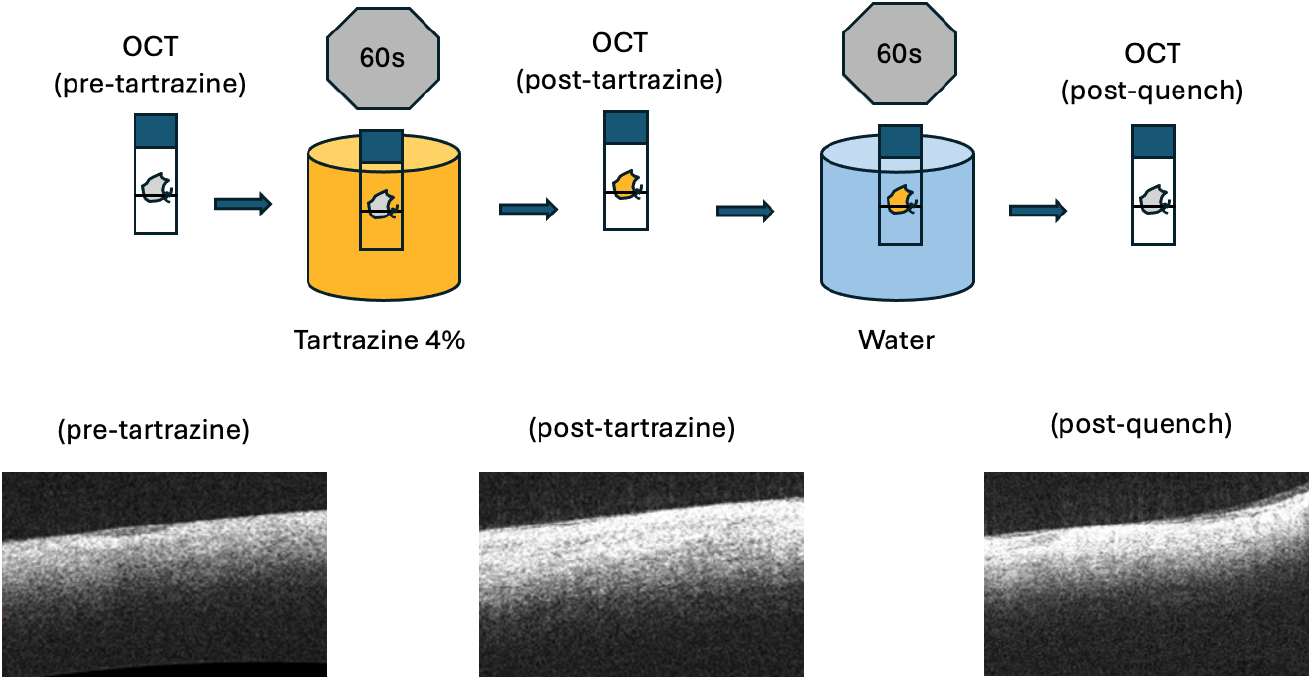
Overview of experimental design. Slide-affixed porcine scleral samples were imaged at three time points: (1) prior to tartrazine immersion, (2) after tartrazine immersion for 60 seconds, and (3) after quenching bath immersion for 60 seconds. Representative OCT images from each of these time points are included underlying the schematic.

After quenching in a water bath for 60 seconds, the penetration depth was reduced again to 354 ± 43.9 µm. All tested samples showed an increase in penetration depth after tartrazine immersion and a decrease in penetration depth after quenching (Figure 2). There was a statistically significant increase in penetration depth after tartrazine exposure (Student’s t = -16.9, p *<* 0.001) and a statistically significant decrease in penetration depth after quenching (Student’s t = 5.7, p = 0.005). Visual inspection of the porcine samples after quenching for 60 seconds demonstrated a reduced orange color to the scleral tissue, but a residual yellow-orange stain remained. As was the case after tartrazine immersion, there were no appreciable changes to the general visual transparency of the samples when inspected with the naked eye. Of note, the samples were simply immersed in the quenching bath and no sample agitation was performed. The OCT penetration of the samples after quenching was found to be greater than that of the samples prior to tartrazine immersion, consistent with the visual appearance of residual dye despite quenching

**Fig. 2.**
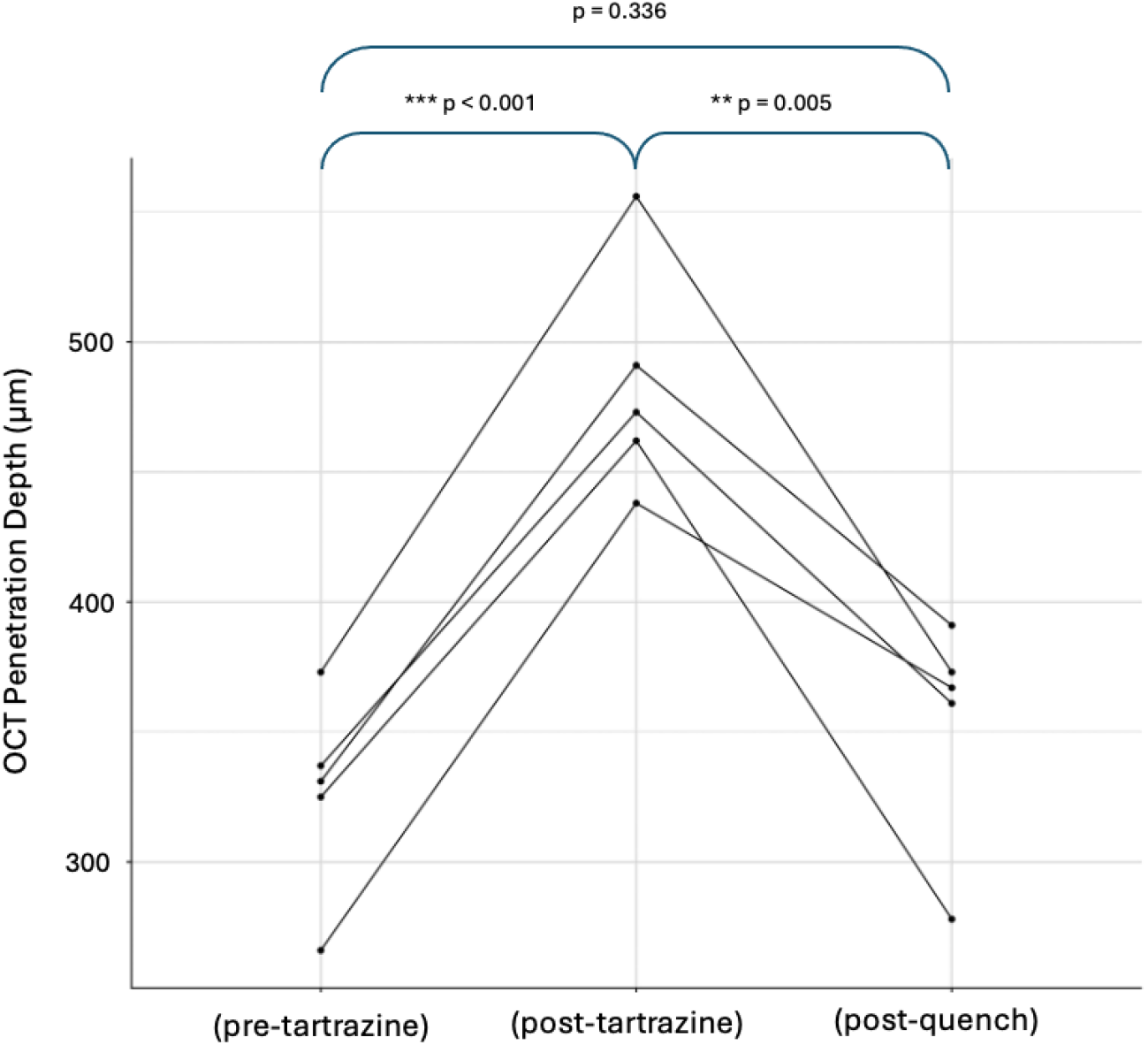
Treatment with 4% tartrazine increases OCT penetration depth in a reversible fashion. Immersion of slide-affixed scleral samples in 4% tartrazine solution for 60 seconds results in measurably increased OCT penetration depth, which is reversed when samples are quenched for 60 seconds in water.

When compared using paired statistical tesitng, however, there was no significant difference between penetration depth prior to sample treatment and after samples completed final quenching (Student’s t = -1.1, p = 0.336) (Figure 2). These data suggest that the quenching process and presumed diffusion of dye particles out of the submerged porcine scleral samples was sufficient to rapidly and repeatably reverse the improved depth of OCT scanning laser penetration.

## 4 Discussion

While the optical clearing effects of porcine sclera noted in our study are of a lesser magnitude than those reported in prior work, ours is the first to show that these effects are present and can be measured even when using an iso-osmolar concentration of tartrazine dye [5, 6]. Furthermore, the dye concentration used in this study, 4%, is much closer to the 3% aqueous solution that was directly tested by Gettings and colleagues for ocular toxicity after direct ocular application [10]. To our knowledge, the Gettings study is the only mammalian in-vivo extended toxicity study of direct ocular surface application of tartrazine dye solutions.

Further, our data suggests that quantifiable clearing effects may be present with extremely limited exposure time, which would improve the potential clinical utility of any tartrazine formulation intended to aid in clinical imaging. The reversal of increased transparency with a very limited time quenching in water that we note during our study suggests that tartrazine readily diffuses out of the scleral tissue in this setting. Complete and prompt removal of dye from the ocular surface is important in reducing the risks of any potential ocular toxicity to patients, and also reduces the risk of any possible cosmetic concerns regarding unattractive yellow-orange staining of the conjunctiva or sclera [13, 14].

It is worth noting that the improvement in OCT imaging depth obtained by isotonic tartrazine was relatively minor as compared to the effects described by extended immersion in high-concentration tartrazine. While prior reports have suggested using transscleral OCT enabled by tartrazine to image suprachoroidal depot administration of medication, it is unlikely that isotonic tartrazine at short exposure times will enable this use case [6]. Despite this limitation, we believe that isotonic tartrazine may instead be of maximal use in imaging structures only partially obscured by the sclera, such as the iridocorneal angle and trabecular meshwork. isotonic tartrazine may also be of use in imaging through partial-thickness anterior lamellar corneal stromal scars.

In addition to imaging, the improved optical penetration for longer-wavelength light afforded by tartrazine may be of therapeutic use as well. For example, the decreased long-wavelength scattering produced by tartrazine application may allow for reduced energy use during transscleral cyclophotocoagulation, and reduced potential for inflammation-associated adverse effects such as choroidal effusion [15, 16].

While reduced dye load to the eye results in a theoretically lower risk of adverse effects, further studies are needed to better characterize the potential ocular toxicity of tartrazine or other dyes intended for in-vivo optical clearing prior to their use in human subjects. Taken together, our data suggests that lowering the concentration and exposure time of tartrazine to levels close to those previously noted to be safe to the ocular surface still results in a measurable improvement in OCT penetration, albeit a more moderate one than than can be achieved with prolonged, high-concentration exposure.

## References

[1] Huang, D., Swanson, E.A., Lin, C.P., Schuman, J.S., Stinson, W.G., Chang, W., Hee, M.R., Flotte, T., Gregory, K., Puliafito, C.A., et al.: Optical coherence tomography. science 254(5035), 1178–1181 (1991)

[2] Drexler, W., Morgner, U., Ghanta, R.K., Kärtner, F.X., Schuman, J.S., Fujimoto, J.G.: Ultrahigh-resolution ophthalmic optical coherence tomography. Nature medicine 7(4), 502–507 (2001)

[3] Zeppieri, M., Marsili, S., Enaholo, E.S., Shuaibu, A.O., Uwagboe, N., Salati, C., Spadea, L., Musa, M.: Optical coherence tomography (oct): a brief look at the uses and technological evolution of ophthalmology. Medicina 59(12), 2114 (2023)

[4] Sardar, D.K., Swanland, G.-Y., Yow, R.M., Thomas, R.J., Tsin, A.T.: Optical properties of ocular tissues in the near infrared region. Lasers in medical science 22(1), 46–52 (2007)

[5] Ou, Z., Duh, Y.-S., Rommelfanger, N.J., Keck, C.H., Jiang, S., Brinson Jr, K., Zhao, S., Schmidt, E.L., Wu, X., Yang, F., et al.: Achieving optical transparency in live animals with absorbing molecules. Science 385(6713), 6869 (2024)

[6] Narawane, A., Trout, R., Viehland, C., Kuo, A.N., Vajzovic, L., Dhalla, A.-H., Toth, C.A.: Optical clearing with tartrazine enables deep transscleral imaging with optical coherence tomography. Journal of biomedical optics 29(12), 120501– 120501 (2024)

[7] Mahmoud, S.S., Elshebley, S.M., Aly, E.M., Awad, S.M., Kamal, G.M.: Second-derivative uv spectral analysis of aqueous humor for eye disease diagnosis and assessing the effects of food additives on ocular health. Scientific Reports 15(1), 17307 (2025)

[8] Barciela, P., Perez-Vazquez, A., Prieto, M.: Azo dyes in the food industry: Features, classification, toxicity, alternatives, and regulation. Food and Chemical Toxicology 178, 113935 (2023)

[9] Feng, J., Cerniglia, C.E., Chen, H.: Toxicological significance of azo dye metabolism by human intestinal microbiota. Frontiers in bioscience (Elite edition) 4, 568 (2012)

[10] Gettings, S., Blaszcak, D., Roddy, M., Curry, A., McEwen Jr, G.: Evaluation of the cumulative (repeated application) eye irritation and corneal staining potential of fd & c yellow no. 5, fd & c blue no. 1 and fd & c blue no. 1 aluminium lake. Food and chemical toxicology 30(12), 1051–1055 (1992)

[11] Team, R.C.: R language definition. Vienna, Austria: R foundation for statistical computing 3(1), 116 (2000)

[12] Wickham, H.: ggplot2. Wiley interdisciplinary reviews: computational statistics 3(2), 180–185 (2011)

[13] Rozman, K.K.: The role of time in toxicology or haber’sc × t product. Toxicology 149(1), 35–42 (2000)

[14] Gaylor, D.W.: The use of haber’s law in standard setting and risk assessment. Toxicology 149(1), 17–19 (2000)

[15] Ishida, K.: Update on results and complications of cyclophotocoagulation. Current Opinion in Ophthalmology 24(2), 102–110 (2013)

[16] Murphy, C., Burnett, C., Spry, P., Broadway, D., Diamond, J.: A two centre study of the dose-response relation for transscleral diode laser cyclophotocoagulation in refractory glaucoma. British Journal of Ophthalmology 87(10), 1252–1257 (2003)

